# Metagenomic characterization of the viral community of the South Scotia Ridge

**DOI:** 10.1101/451732

**Authors:** Qingwei Yang, Chen Gao, Yong Jiang, Min Wang, Xinhao Zhou, Hongbing Shao, Zheng Gong, Andrew McMinn

## Abstract

Viruses are the most abundant biological entities in aquatic ecosystems and harbor an enormous genetic diversity. While their great influence on the marine ecosystems is widely acknowledged, current information about their diversity remains scarce. Aviral metagenomic analysis of two surfaces and one bottom water sample was conducted from sites on the South Scotia Ridge (SSR) near the Antarctic Peninsula, during the austral summer 2016. The taxonomic composition and diversity of the viral communities were investigated and a functional assessment of the sequences was determined. Phylotypic analysis showed that most viruses belonging to the order *Caudovirales,* in particular, the family *Podoviridae* (41.92-48.7%), which is similar to the viral communities from the Pacific Ocean. Functional analysis revealed a relatively high frequency of phage-associated and metabolism genes. Phylogenetic analyses of phage TerL and Capsid_NCLDV (nucleocytoplasmic large DNA viruses) marker genes indicated that many of the sequences associated with *Caudovirales* and NCLDV were novel and distinct from known complete phage genomes. High *Phaeocystis globosa* virus virophage (Pgvv) signatures were found in SSR area and complete and partial Pgvv-like were obtained which may have an influence on host-virus interactions in the area during summer. Our study expands the existing knowledge of viral communities and their diversities from the Antarctic region and provides basic data for further exploring polar microbiomes.

**Importance:** In this study, we used high-throughput sequencing and bioinformatics analysis to analyze the viral community structure and biodiversity of SSR in the open sea near the Antarctic Peninsula. The results showed that the SSR viromes are novel, oceanic-related viromes and a high proportion of sequence reads was classified as unknown. Among known virus counterparts, members of the order *Caudovirales* were most abundant which is consistent with viromes from the Pacific Ocean. In addition, phylogenetic analyses based on the viral marker genes (TerL and MCP) illustrate the high diversity among *Caudovirales* and NCLDV. Combining deep sequencing and a random subsampling assembly approach, a new Pgvv-like group was also found in this region, which may a signification factor regulating virus-host interactions.

## Introduction

Viruses exist wherever other life is found, including in the deep ocean and polar areas. Arguably, viruses are by far the most numerous, genetically diverse, and pervasive biological entities in the aquatic ecosystem(1, 2). They are critical mortality agents of both eukaryotes and prokaryotes, affecting the abundance and diversity of microbial communities as well as global biogeochemical processes and energy fluxes by causing lysis of a large proportion of both autotrophic and heterotrophic prokaryotes, shunting nutrients between particulate and dissolved phases(3-8) and modifying the efficiency of the carbon pump(9). Furthermore, bacteria and protists as vehicles for viral reproduction, and their genetic diversity was shaped by virus-mediated horizontal gene transfer, allowing viral genes to spread far and wide(2, 10).

The ecology of prokaryotes and protists, especially Antarctic phytoplankton during summer(11-16), together with the major role of viruses in prokaryotes and eukaryotic phytoplankton mortality(17-20) have been well studied. However, due to the geographical location and the difficulty of culturing viral hosts, an understanding of virus diversity and viral community structures in coastal regions of Antarctica including Sub-Antarctic areas is still lacking. So far, there have been few studies, based on culture-independent methods such as metagenomics and single-cell genomics, were conducted to analyze the DNA and RNA viral communities in Antarctic environments including freshwater habitats(21-25), the Southern Ocean closed to Western Antarctic Peninsula(26), sediment as well as soil(27, 28). These studies all pointed toward a high viral biodiversity in these Antarctic ecosystems. However, despite the virome diversity information derived from these special habitats in Antarctic, there have been few studies from the open sea near the Antarctic Peninsula.

In this study, we conduct an analysis of three viromes from South Scotia Ridge (SSR) seawater samples including two from the surface and one from the bottom (water depth=500 m) in an area influenced by Antarctic Circumpolar Current flow (ACC)(29). The major steps of the analysis were: (i) the determination of the composition of these three viromes and the dominant viral species, (ii) a comparison of these viromes with those from different habitats, (iii) the potential functional analysis, (iv) a phylogenetic and/or genomic analysis of the major viral groups that were present in these viromes.

## Results

### Overview of SSR viromes

The metagenome of three seawater samples collected from the SSR in Antarctic Peninsula region, ranged in temperature from −0.04 to −0.57 °C and in salinity from 34.37%o to 34.57%o (Fig. S1, Table S1). After extraction a total of 129,710,606 paired-end 150 bp sequences, with 109,923,264(84.75%) reads passing the quality screening (Table 1), were obtained.

**Table 1.** The remaining number of quality-controlled reads and nonredundancy contigs

| Process |  | SSR viromes |  |  |
| --- | --- | --- | --- | --- |
|  |  | D39s | DA4s | DA4b |
| Quality control | Raw reads | 42547324 | 43791908 | 43371374 |
|  | Cut adaptor | 40670620(95.59%) | 41883752(95.64%) | 41353540(95.35%) |
|  | Q20>20% | 35357306(83.10%) | 37567224(85.79%) | 38397082(88.53%) |
|  | Q30>30% | 35334370(83.05%) | 36385270(83.09%) | 38203624(88.08%) |
| Assemble | All assembled contigs(>500bp) | 2418081 | 3699559 | 1693019 |
|  | Non-redundancy contigs | 145023 | 135910 | 234648 |
| Mapping | Mapped reads | 15307526(43.32%) | 12760519(35.07%) | 21197289(55.49%) |

Best BLAST Hit (E-value < 1e^−3^) affiliations of unassembled high-quality reads from the three data sets are consistent with viral metagenomes analyses published so far, as more than three-quarters of the reads (75.7-88.24%) did not show any significant sequence similarity to current NCBI nr data (Fig. 1a). According to the NCBI nr and viral RefSeq annotation, 3.31-10.87% and 2.68-6.61% of the reads were classified as viruses respectively (Fig. 1b). Acomparison of the annotation results of NCBI nr and viral RefSeq, found that the virus sequences annotated with virus in the NCBI nr database were more abundant than those in viral RefSeq (Fig. 1c), indicating that there is a number of sequences belonging to an unidentified virus that viral RefSeq excluded, such as uncultured Mediterranean phage uvMED.

**Fig. 1.**
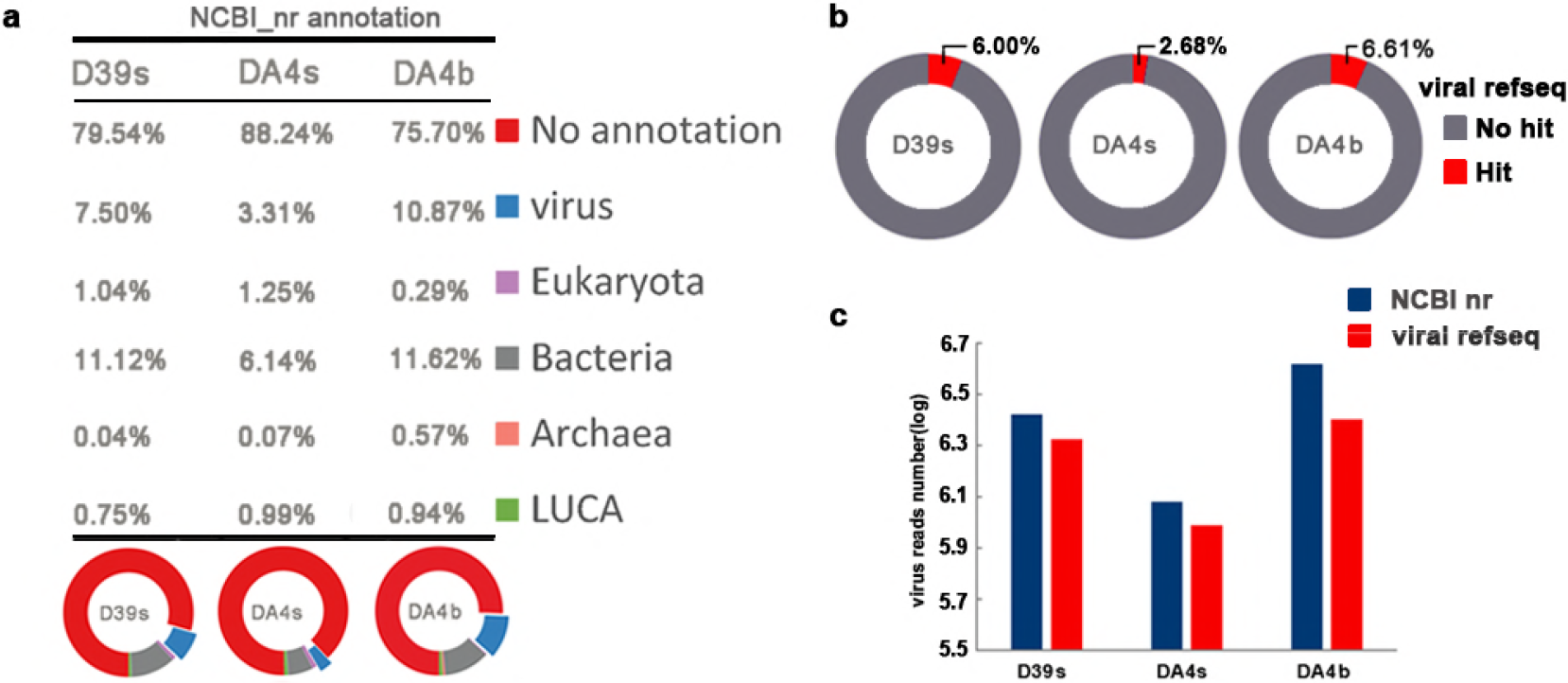
Taxonomic assignment of metagenomic reads (a)percentage of the sequence reads classified by the taxonomic grouping based on BLASTX similarity search with NCBI nr database (E-value<1e^−3^). Sequences with no hits with E-value > 1e-3 were regarded as unidentified reads (“no annotation” category in the table and red in the pie graphs). LUCA (green) denotes reads that could not unambiguously be assigned to a domain of life. (b) taxonomic assignment of metagenomic reads based on BLASTX similarity search with viral RefSeq database (E-value<1e-3). (c) compare the annotation results of NCBI nr and viral RefSeq.

### Taxonomic Diversity Analysis

The BLAST data results (against viral RefSeq) of the virome composition are visualized using the Krona tool (Fig. S2, Fig. S3 and Fig. S4) (30)and show that, as excepted, the majority of viral reads (93.69-95.16%) with significant hits belonged to double-stranded DNA (dsDNA) viruses with no RNA stage. These were largely comprised of members of the *Caudovirales* comprising the families *Podoviridae, Siphoviridae* and*Myoviridae,* with similarities to single-stranded DNA (ssDNA)viruses and RNA viruses were also observed (Table 2 and Table 3). *Podoviridae*sequences (41.92%-48.7%) were the most abundant in all three viromes followed by *Myoviridae*(22.92-29.46%) and *Siphoviridae* (11.92-14.08%). Viruses from the *Phycodnaviridae* (infecting algae) and*Mimiviridae* (infecting amoebas and algae) were more abundant in surface waters than bottom water (D39s: 3.57% and 0.16%; DA4s: 2.22% and 0.22%; DA4b: 1.32% and 0.10%, respectively). There was a significant proportion of virophages that prey on phycodnaviruses in surface water, approximately 2% in D39s (Table 2). The top ten most abundant viral species (Fig. S5) including *Puniceispirillum* phage HMO-2011(31) *(Podoviridae,* circular genome), a phage infecting a bacterium of the SAR116 clade, was the most abundant in the SSR virome (18.50-25.75%), nearly accounting for 25% in DA4s, and Pelagibacter phage HTVC008M (32) *(Myoviridae,* linear) aT4-like myovirus infecting a SAR11 bacteriophage was second most abundance (8.6-11.11%).

**Table 2.**
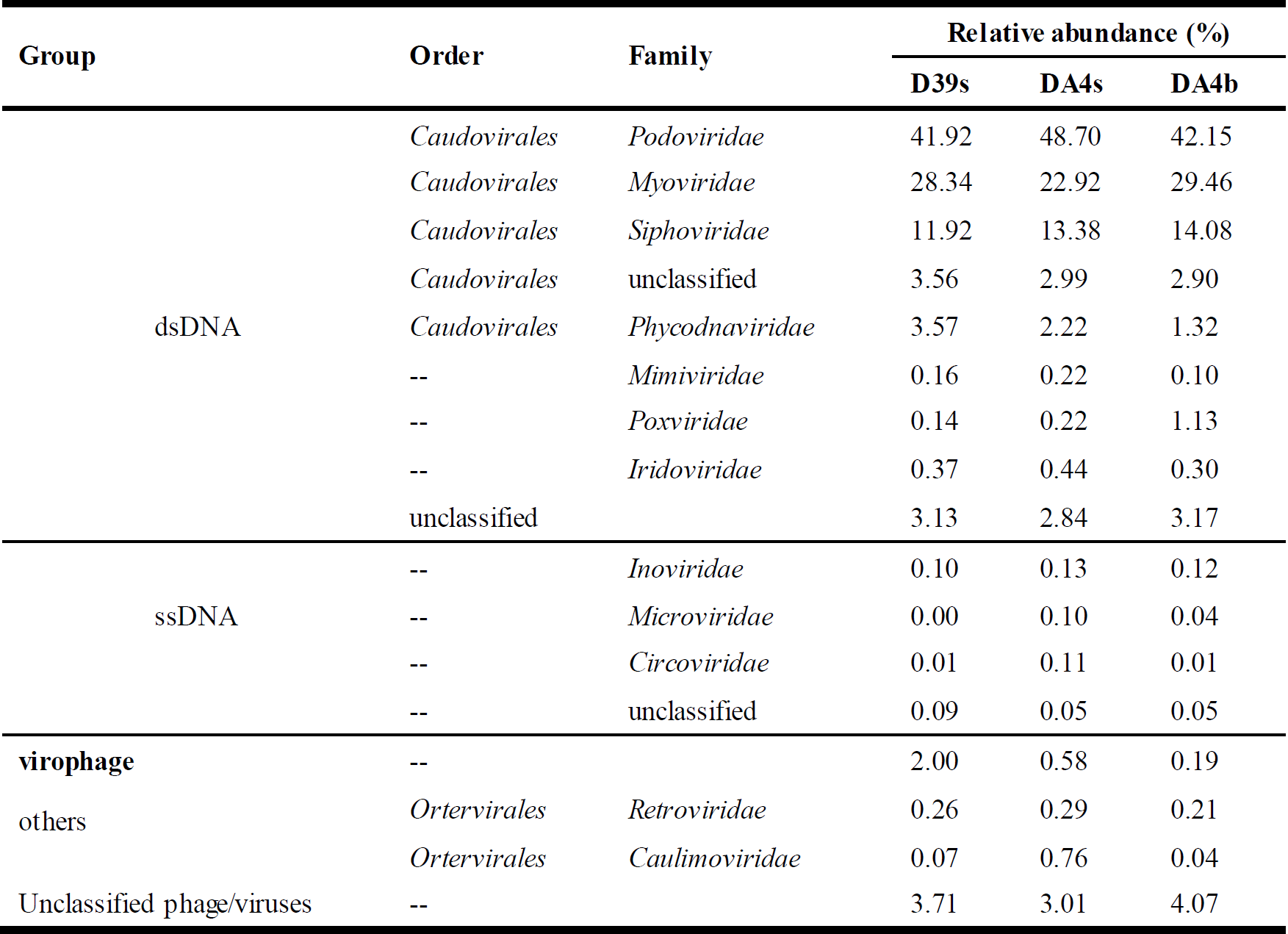
Classification of reads from viromes hitting viral sequences

**Table 3.**
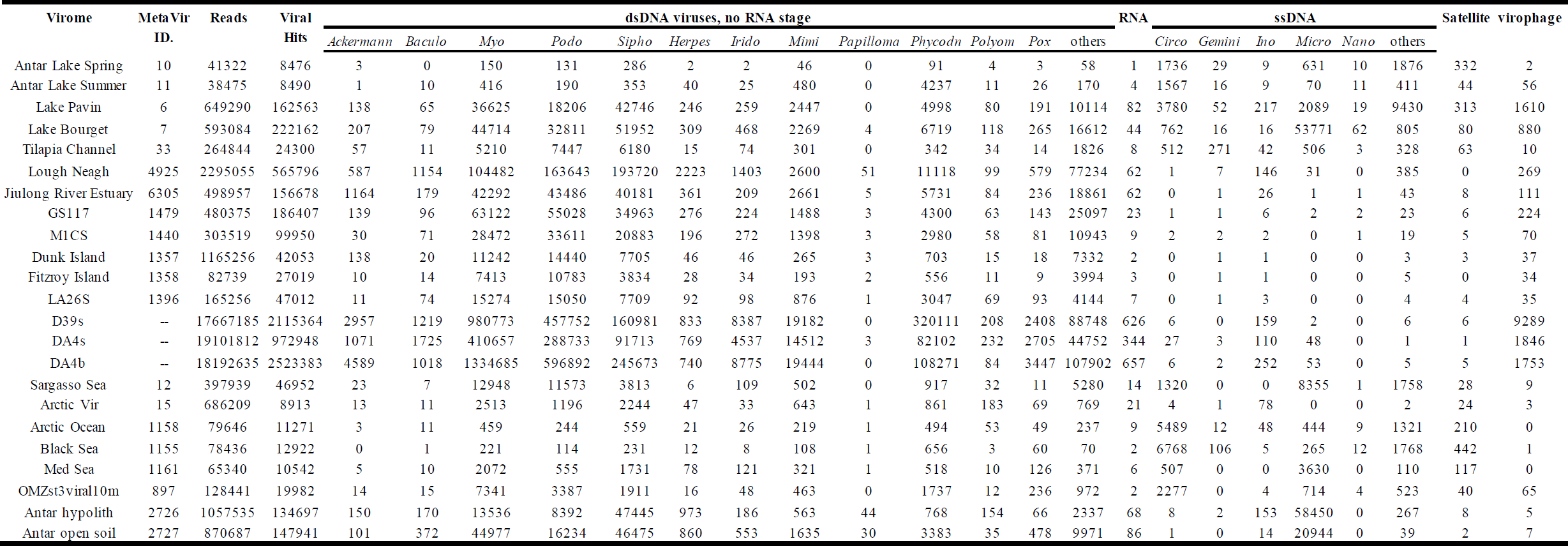
Distribution of sequences from the SSR viromes and twenty previously published viromes as determined by the indicated BLAST comparison to viral Refseq(E-value<1e-3)

### Comparison with other Virome

To compare viromes between present study and previously published data sets, twenty viromes from different habitats were selected from MetaVir (see Materials and Methods for details). The result showed that the three SSR viromes were most closely related to ocean surface samples from the previous studies, except for the samples from ETSP-OMZ and SAR (Fig. 2), from which statistically significant differences were measured (p < 0.001). At the ocean surface, virome taxonomic at the family level was dominated by the *Caudovirales (Myoviridae, Siphoviridae, Podoviridae),* which collectively contributed 43.74-92.03% of the genomes. Viromes within special habitats including deep-ocean surface sediments, ETSP-OMZs, Antarctic freshwater, soil and hypolith are dominated by *Ciroviridae* and *Microviridae* members of ssDNA viruses which contributed 25.45-88.45% (Fig. 3, Table 3). Less than 5% of these viromes sequences showed any similarity (E-value < 1e^−3^) to the SSR viromes (Table 4).

**Table 4.**
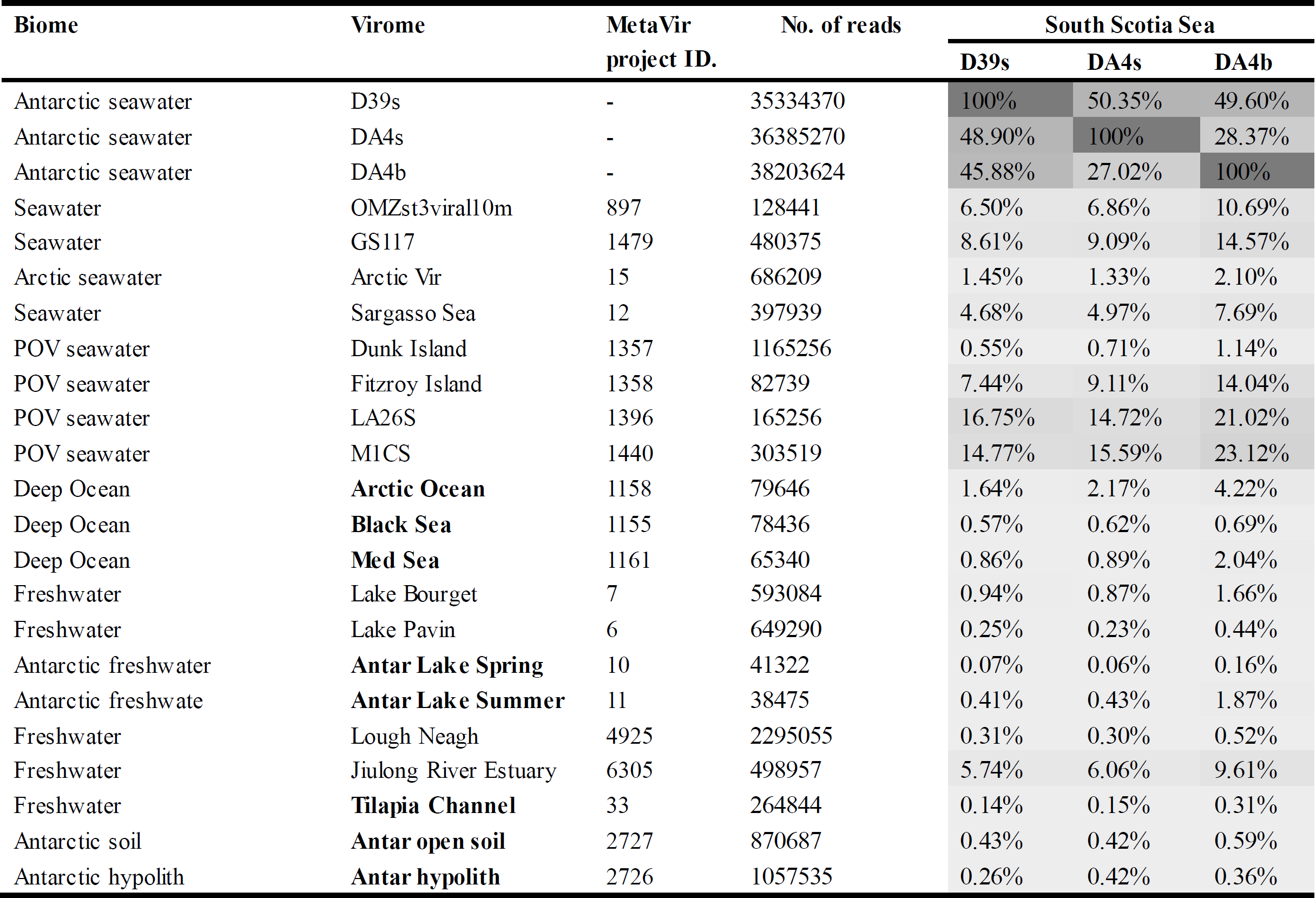
The table shows the percentage of reads in other published viromes obtained from MetaVir with a significant similarity (BLASTN, E-value<1e^−3^) to the SSR viromes

**Fig. 2.**
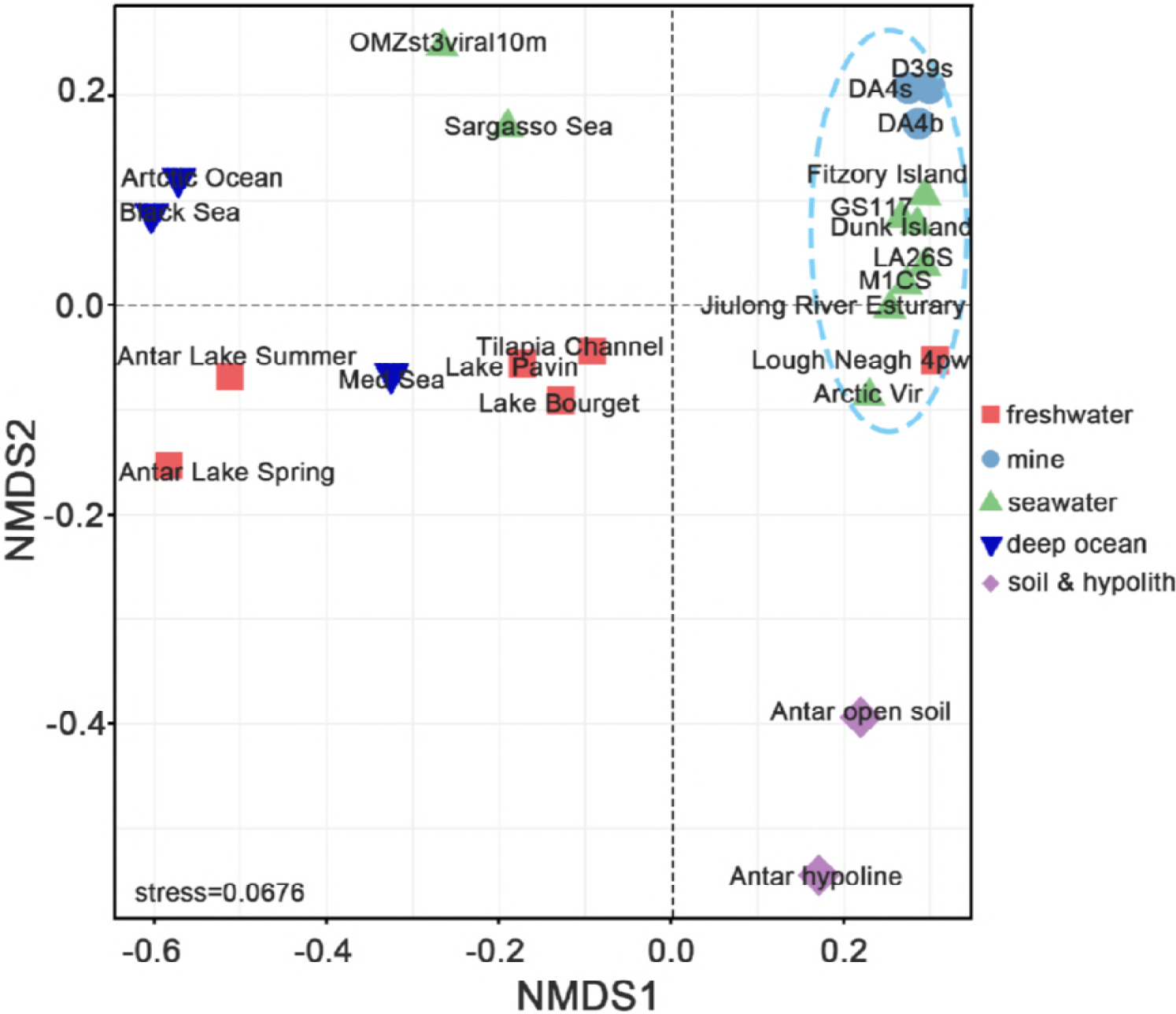
Comparison viromes between SSR area and other environmental viromes depending on taxonomic composition. Twenty environmental viromes were available on MetaVir2, obtained from different habitats including freshwater, seawater, deep-sea surface sediments, soil and hypoline.

**Fig. 3.**
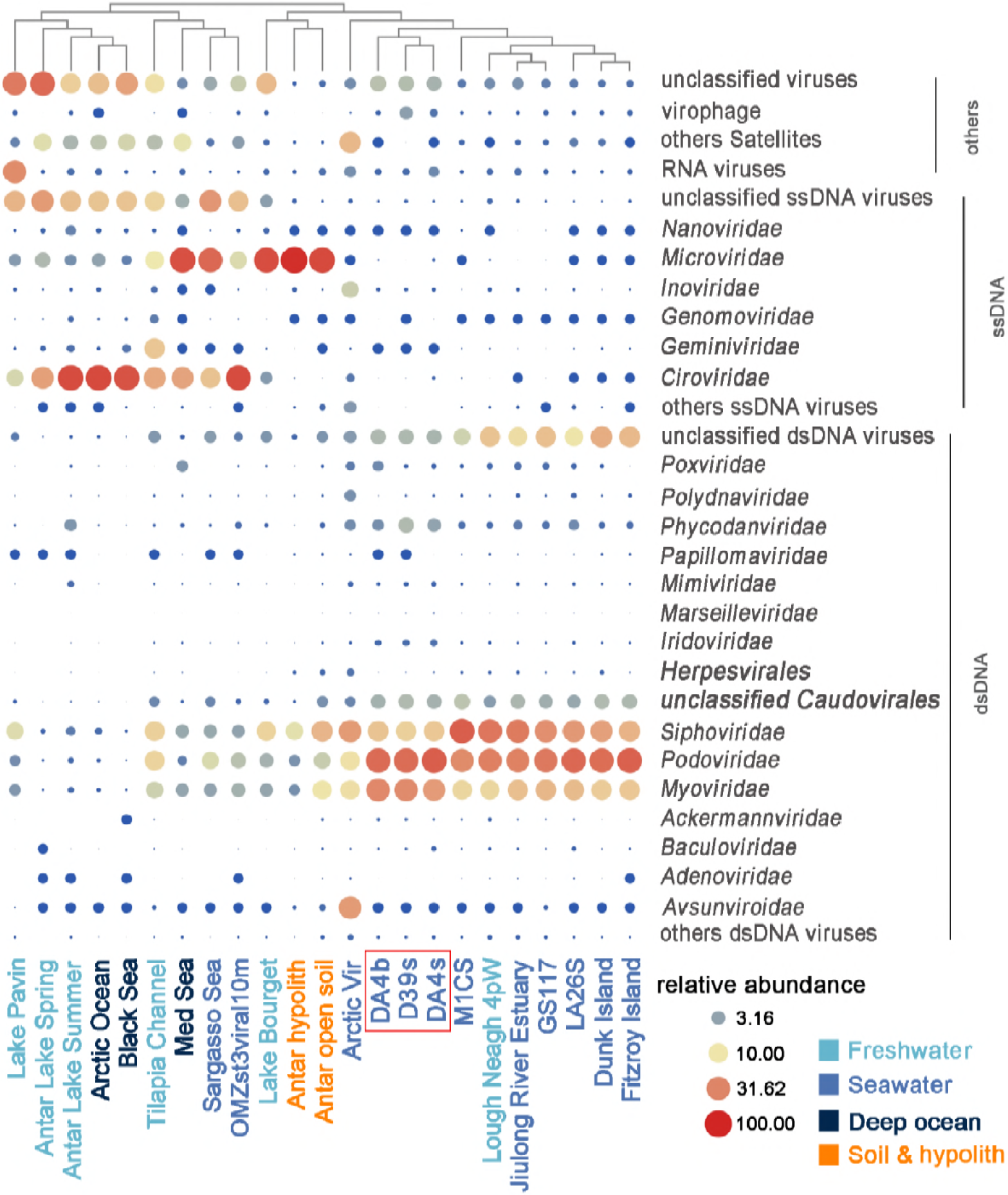
The relative abundance of viral sequences (normalized with genome length) largely at the family level in each different habitat virome. Points size indicate the value of relative abundance.

### Contigs and function analysis

As the contigs assembled by the random subsampling approach could still contain redundant sequences derived from the same (or closely related) populations contigs derived from the same population were merged into clusters with 90% global average nucleotide identity by cd-hit-est, resulting in 145,023(D39s), 135,910(DA4s) and 234,648(DA4b) non-redundant genome fragments (>500 bp) respectively (Table 1). Of these 43.32%(D39s), 35.07%(DA4s) and 55.49%(DA4b) quality-filtered reads were assigned to nr contigs. The putative functions of the annotated ORFs from the nr contigs dataset were predicted using MG-RAST, which assigns sequences to metabolic categories based on their Best BLAST Hit against the SEED database (E-value < 1e^−5^). Using the subsystems approach, nearly 25% (17.54-26.46%) of the annotated proteins fell into the ‘Phage, Prophage, Transposable elements or Plasmids’ (Fig. 4). Phage structural, integration/excision and DNA metabolism-related proteins were most commonly identified and 10-11.96% of them classified into ‘Clustering-based subsystems’ with phage endolysin commonly found in this category. The other SEED functional annotation categories showed the metabolism of amino acids, carbohydrates, cofactors, vitamins, proteins, RNA, DNA, and nucleosides/nucleotides were the dominant annotations. In these categories, many proteins could be phage-related (or possible cellular origin) such as DNA polymerases and helicase. These hits were also found in the Pfam and COGs databases (see IMG system), with ‘replication, recombination and repair’ being the most common protein categories identified.

**Fig. 4.**
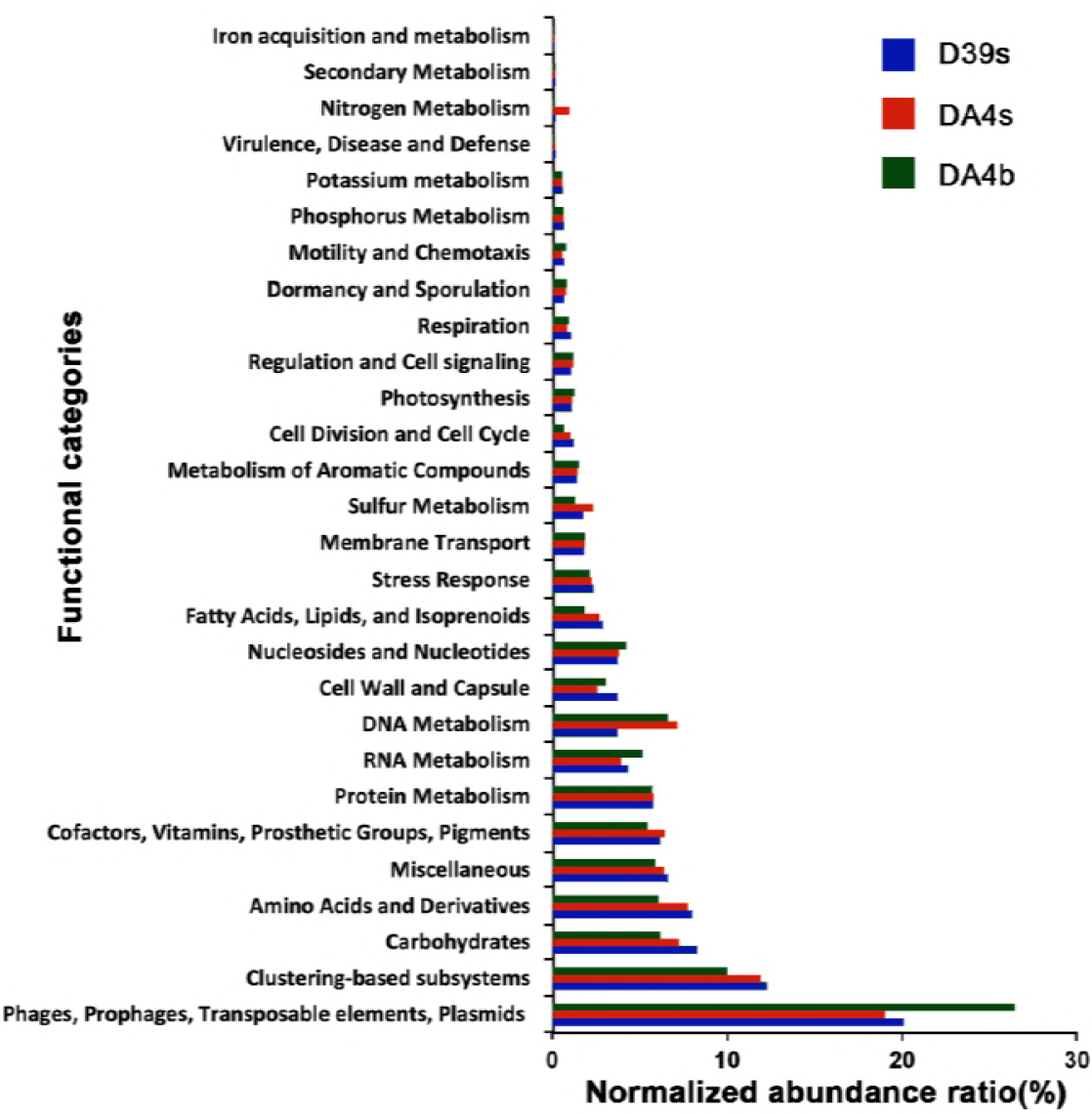
Composition of predicted functional genes of the SSR contigs. The CDSs were compared with the SEED database using subsystems in MG-RAST. The metabolic categorization is based on the sequences Best BLAST Hits in the SEED database curated subsystems (E-value <10^−5^).

### Phylogenetic tree analysis

#### Terminase phylogeny

An ML phylogenetic analysis of the phage large terminase subunits identified in this work is shown in figure 5. The topology of the phylogenetic tree clearly shows that the majority of the SSR viromes’ TerL amino acid sequences were widely distributed among the *Myo-, Sipho-* and *Podoviridae,* most of which were phylogenetically distant to known complete phage genomes (deposited in the NCBI database), and six new groups (in blue) did not cluster with any known species. This is supported by high bootstrap values, highlighting but important uncharacterized diversity for *Caudovirales* in SSR.

**Fig. 5.**
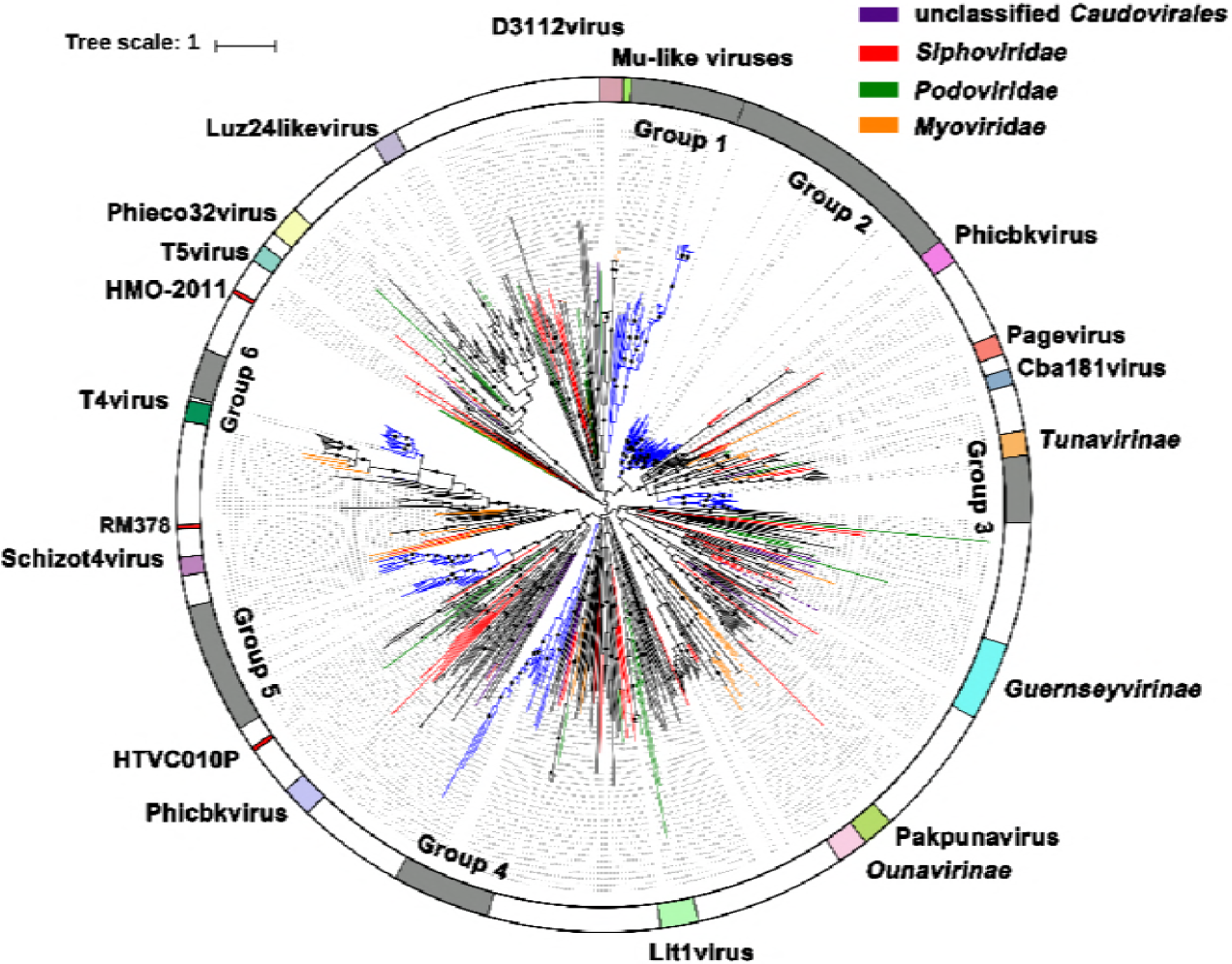
Terminase phylogeny. A maximum-likelihood phylogenetic tree of *Caudovirales* terminase large-subunit domains (PF03237) is shown (1000 iterations, JTT+G model). Only bootstrap values of > 50% are indicated at the nodes of the tree and bootstrap scores greater than 90% are indic ated with a black dot. Average branch length distance of leaves less than 0.4 were collapsed and shown as triangles. Reference sequences are marked (see color legend at the top) and new groups are highlighted in blue.

#### Capsid_NCLDV phylogeny

ML phylogenetic tree, based on the MCP including a group of putative MCP of Pgvv-like infected *Phaeocystis globosa* virus (Pgv), is shown here (Fig. 6). The MCP tree shows that several sequences from SSR viromes are closely related to known NCLDV, mainly those belonging to Phycodnaviruses, and these can be classified into Prasinovirus, Pgv, and Pgvv. The three clades differed from Phycodnaviruses and Mimivirus MCPs, forming three distinct groups with the well-supported clades; one of the clades, marked as Group3, was only found in the surface ocean of DA4 station. The Pgv group, included five new Pgvv-like MCPs, was distantly related to the Pgv group and had a higher relative abundance at the surface than at the bottom.

**Fig. 6.**
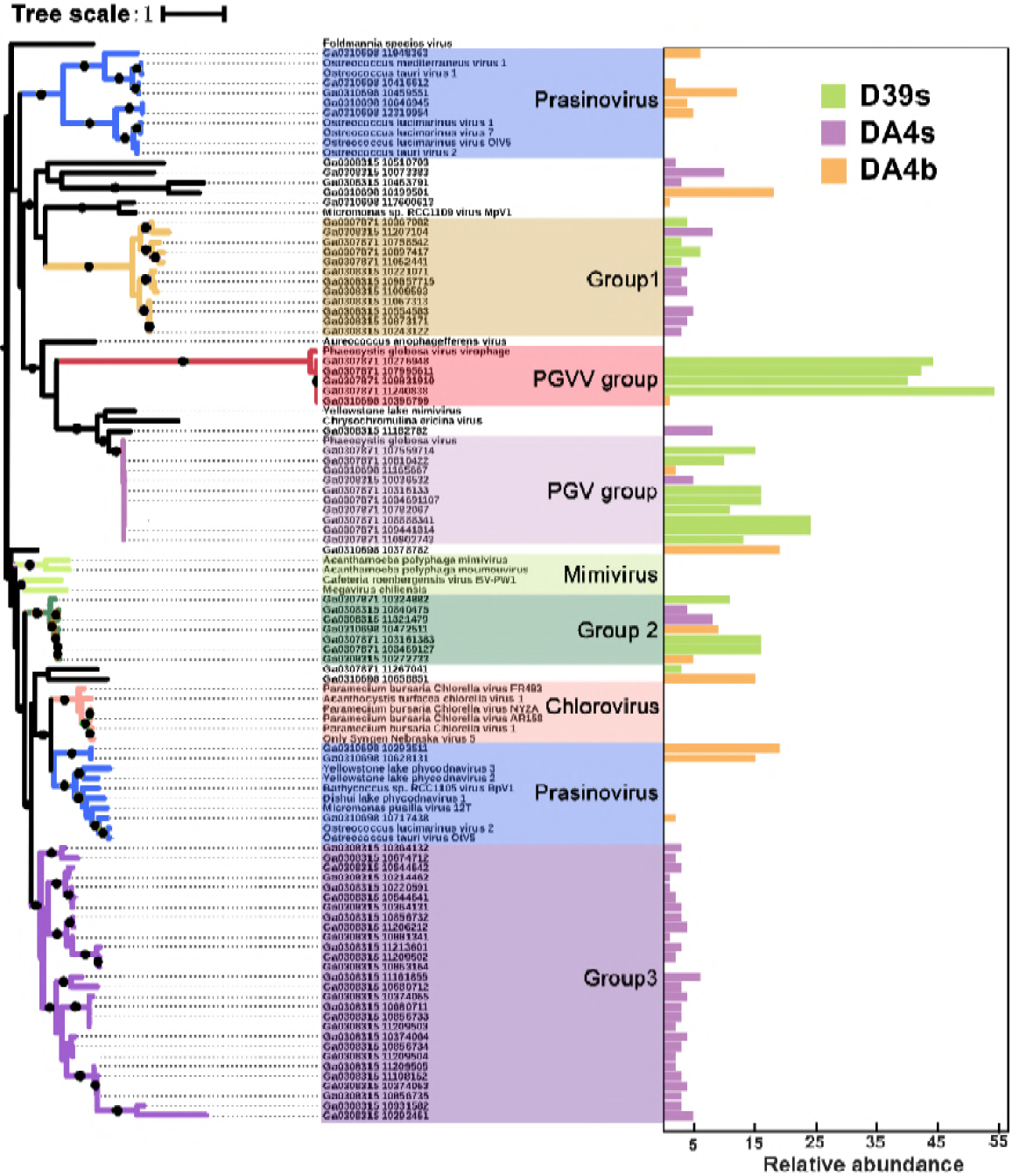
Capsid_NCLDV phylogeny. A maximum-likelihood phylogenetic tree drawn from the capsid_NCLDV (PF04451) and six virophage putative major capsid (MCP) protein multiple alignment is shown (1000 iterations, JTT+G model). Bootstrap scores greater than 90% are marked with black dots. Each MCP is associated with an abundance profile (right) that displays the relative abundance of the contig across the three SSR viromes (based on normalized coverage).

### Novel Pgvv group

From the MCP phylogenetic tree one distinct group of virophage was defined for which there is one known related virophage genome. However, it was still very different from the known Pgvv. An alignment of Pgvv-like group genomes is shown in figure 7. The Pgvv-like group genomes appear to have a relatively high (37.36-38.17%) GC content expect for the GC content of Pgvv-like 04 genome (GC, 35.85%) was similar to Pgvv (GC, 35.8%). All virophages share four homologous proteins or domains: 1) packaging ATPase (ATPase), 2) lipase, 3) major capsid protein (MCP), 4) minor capsid protein (mCP). In addition, Pgvv-like 02 also contain the OLV11-like tyrosine recombinase (Yrec) gene which is only distantly related to the OLV11-like family(33). Three genes with functional annotation (in yellow), absent in the Pgvv genome, were carried by Pgvv-like sequences, including putative primase-helicase and DNA methyltransferase genes in Pgvv-like 02 and recombination endonuclease VII gene in Pgvv-like 04. These further indicate that these viruses maybe belong to a new Pgvv-like group.

**Fig. 7.**
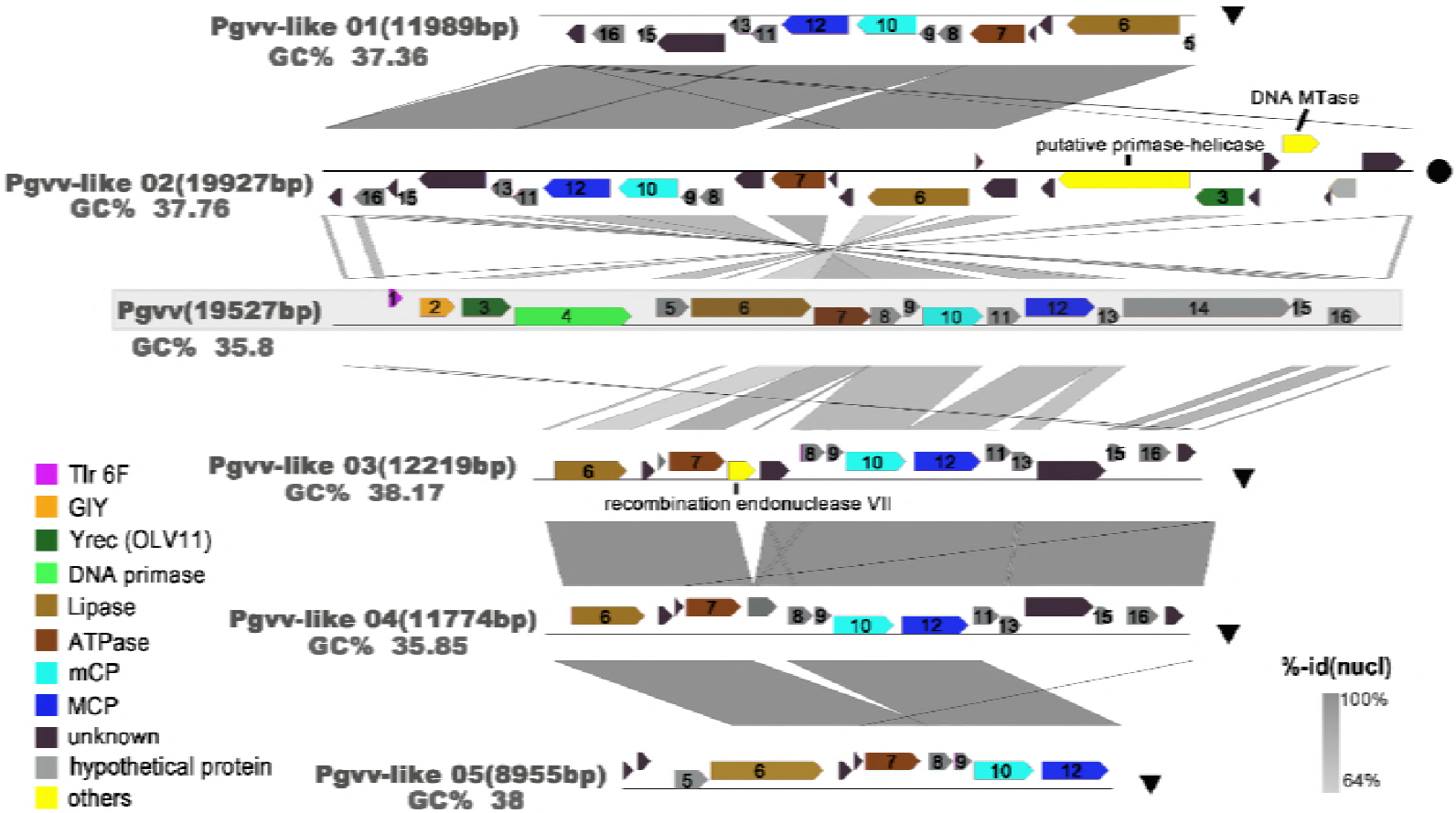
Overview of genomic synteny and similarities between Pgvv-like group. The complete Pgvv reference was covered by grey shadow. A color scale for percent identity (nucleic) is shown at the bottom right. The name, percent GC content (GC%), and length for each genome are also indicated. Genes are colored according to their functional affiliation. Tlr 6F, Toll-like receptor 6 family; GIY GIY-YIG family nuclease; MCP, major capsid protein; mCP, minor capsid protein; Yrec (OLV11), OLV11-like tyrosine recombinase.

## Discussion

Marine viral communities are still largely undescribed and many basic issues, such as their global ocean distribution and their actual genetic and species richness, remain unanswered(2, 9, 34). With the advent of metagenomic methods, associated with high-depth sequencing and meta-analyses of bioinformatics, an increasing number of studies have been conducted(35, 36). So far, only a few of these have focused on viral communities from the Antarctic region and most of these are from special habitats, such as freshwater lakes, hydrothermal vents and soils. For example, high diversity viral communities have been described from a lake (21), soils(28), and hydrothermal vents sites(37). As regards the sea water viromes, the important role of temperate viruses (that is, those capable of both lysogeny and lytic replication) in western Antarctic Peninsula dsDNA viral communities has recently been revealed(26) and differences in viral community composition between the subtropical Indian and Southern Oceans has been identified(38). The number of reads identified as either bacteria or eukaryote was similar to that reported from viral metagenomes of other environments(34, 39), indicating a certain degree of bacterial and eukaryotic contamination of the metagenomes. In addition, the high proportion of no annotation sequences and relatively low number of rRNA and tRNA genes (< 1%) matching sequences (Table S2) indicates bacterial and eukaryotic contamination, previously reported to occur with TFF-based concentration methods(40). Because bacterial genes can be packaged into generalized transduced phage particles(41, 42), the bacterial-like sequences might have come from excised prophages mistakenly annotated as bacterial and/or from genes of bacterial origins that were transferred to their phages(39).

BLAST searches showed that >75% of the sequences before assembly did not have homologs in current sequence databases. This result is consistent with the results of the previously published viral metagenomic projects(38, 43-45) and such a high proportion of unknown sequences reemphasizes that most of the biodiversity in the viral world is still undiscovered. The SSR viromes were largely dominated by *Caudovirales,* including Myoviruses, Siphoviruses, and Podoviruses, which are the dominant viral type recovered during metagenomic analyses of marine environments(2, 43). In the three SSR viromes investigated here, the largest number of reads (>40%) were related to Podoviruses and ca. 13% of reads were of Siphoviruses (viruses which infect photosynthetic bacteria such as *Prochlorococcus* and *Synechococcus)* (in bold, Table S3). Consistent with previous investigation (31, 46), *Puniceispirillum* phage HMO-2011, that infects “*Candidatus Puniceispirillum marinum*” strain IMCC1322 of the SAR116 clade, and the *Pelagibacter* phage group (HTVC008M, HTVC010P, HTVC011P and HTVC019P), infects SAR11 populations were widespread and most abundant in SSR. Both of SAR11 and SAR 116 clades play important roles in oceanic Dimethyl sulfide (DMS) production and biogeochemical sulfur cycles, especially via bacteria-mediated dimethylsulfoniopropionate (DMSP) degradation(47, 48). Interestingly, Pgv is the tenth most abundant viral species in SSR region (2.53% in D93s), infecting the temperate algal species *Phaeocystis globosa(49).* In the Antarctic, however, one of the most abundant *Phaeocystis* species is *P. antarctica* (50) and the *P. antarctica*virus has still not been isolated and identified, which may indicate the high genome similarity between *P. antarctica* virus and Pgv. Despite being in a cold marine environment with an average temperature below 0 °C, the SSR viral community had a structure similar to those found in the Pacific Ocean. However, there were still significant differences in nucleic acid levels and it is likely that the genotype of many viruses changed to allowing them to infect psychrophiles and thus evolve into new viral groups. The previously studied viromes from deep-ocean surface sediments, ETSP-OMZs, Antarctic freshwater, soil and hypolith, in which ssDNA viruses played dominated roles, were clearly different from those of the SSR. However, all of those viromes (except those from the deep-ocean surface sediment) were amplified using multiple displacement amplification (MDA) with phi29 polymerase, so the genomes of the ssDNA viruses could have been selectively amplified (51, 52), which may have led to an overestimation of the role of ssDNA viruses. Although existence bias from MDA in these studies and the prevalence of *Caudovirales* sequences has been observed in most marine viromes, previously published research on global morphological analysis of marine viruses, conducted by the Tara Oceans Expedition, showed that non-tailed viruses (largely ssDNA and RNA) numerically dominate the upper oceans (53)and small, non-tailed viruses were undoubtedly underestimated in SSR region also.

The deep sequencing method, combined with a random subsampling assembly approach, made it possible to obtain a nearly complete viral genome and create phylogenetic analyses on marker genes. Analysis of the major viral groups found in the SSR viromes showed a very broad diversity and many previously unknow virotypes. The terminase gene which is responsible for DNA recognition and initiation of DNA packaging, is an essential component of all head-tail phages *(Caudovirales),* encoding the molecular movements that translocate DNA into empty capsids (54). There is a large diversity of terminases, which can be used to resolve different Caudoviruses groups(55). The NCLDV comprises a monophyletic group of viruses infecting both animals and a diverse range of unicellular eukaryotes, including the *Phycodna-, Mimi-, Asco-, Asfar-, Irido-* and *Poxviridae* families. The MCP of NCLDV (capsid_NCLDV), a redox protein that encodes complex DNA replication and transcription systems and involved in the formation of disulfide bond in virion membrane proteins, is relatively conservative among NCLDVs evolution(56-58). Using phylogenetic trees based on these two viral marker genes (TerL and MCP), illustrated the high diversity among *Caudovirales* and NCLDV A high proportion of TerL sequences were distributed both far from the reference and also far from each other, highlighting both the richness of *Caudovirales* in the SSR communities and also the absence of closely-related reference sequences. In addition, some SSR virome sequences appear to form a new clade (Group 6) related to the T4 viruses, one of the best described *Caudovirales* families.

The topology of the MCP tree and genomic comparisons strongly suggest that the five putative virophage genomes are more closely related to the Pgvv than to other NCDLV families, including the Pgvv host. The Pgvv-like group also has a high relative abundance. The Lotka-Wterra simulation demonstrated that virophages promote secondary production through the microbial loop by reducing overall mortality of the algal cell after a bloom and increasing the frequency of blooms during the summer(18). According to the above model, it can be inferred that the Pgvv-like group plays a previously unrecognized role in regulating virus-host interactions in SSR area during summer.

## Methods and materials

### Sample Collection and Sequencing

Seawater samples, including two surface waters and one bottom water (Table S1), were collected during the austral summer (December 2016) from two sites (D39 close to the edge of the Powell Basin and DA4 near to Clarence and Elephant Islands, Fig. S1) on the southern flank of SSR situated between South America and Antarctica Peninsula. Seawater temperature and salinity were recorded with a CTD profiler (SBE9/11 plus V5.2, Sea-Bird Inc., USA). Water for biological and chemical analysis was collected with Niskin bottles attached to the CTD profiler and was prefiltered with a 20 μm mesh to remove large particles.

The virome samples were processed immediately according to Sun et al(59). Briefly, the samples were sequentially filtered through 3 μm and 0.22 μm pore size filters to remove any microorganisms, and then a Two-step tangential flow filtration (TFF) with 50-kDa cartridge (Millipore, MA, USA) was used to concentrate the viruses to a final volume of ca. 50 ml and stored at −80 °C. The samples were further concentrated by Polyethylene glycol (PEG-8000) precipitation (10% w/v) and incubated at 4 °C overnight. The concentrate was then centrifuged at 8000 g for 80 min at 4 °C and suspended in 200 μl SM buffer. Finally, DNA was extracted using the phenol/chloroform/isoamyl method and precipitated with ethanol without random amplification. High-throughput sequencing was performed by Novogene (Beijing, China) using Illumina Hiseq X ten (Paired End sequencing, 2μ150 bp).

### Virome Composition Analysis

A series of quality-screenings were undertaken to further remove low quality reads; these followed the quality control protocols of the sequencing company. First, adaptor sequences were trimmed by cutadapt (v 1.14). Subsequently, sequences that had a Phred score of at least 20 with a minimum length of 100 bp and had a Phred score of at least 30 with a minimum length of 88 bp were removed from libraries.

In order to avoid chimeras, SSR virome sequences were analyzed without assembly and queried by Diamond (60)against the NCBI non-redundant(nr) protein database(ftp://ftp.ncbi.nlm.nih.gov/blast/db/FASTA/) and RefSeq complete viral genomes protein references (viral RefSeq) database(ftp://ftp.ncbi.nlm.nih.gov/refseq/release/viral/), setting a maximum E-value of 1e^−3^. Taxonomic identification was assigned based on best similarities and the relative taxonomic was normalized against complete viral genome length.

### Virome Composition Analysis

Twenty previously published viromes taxonomic compositions, with the same maximum E-value based on reads number were selected from MetaVir to compare with this study(61). These were obtained from a variety of habitats, including six temperate freshwater lakes(Lake Bourget and Lake Pavin(62), Antarctic lakes (21), Lough Neagh(63), Tilapia Channel(64)), nine seawater sites from eastern tropical South Pacific Oxygen minimum zones (ETSP-OMZ)(65),the Indian Ocean(66),the high salinity Jiulong River Estuary (67), Dunk Island, Fitzroy Island, LA26S and M1CS of POV(44), the Arctic Ocean and Sargasso Sea (SAR)(68)), three deep-sea surface sediments (Arctic Ocean, Black Sea and Mediterranean sea(69), soil and hypolith(70)) (Table 3). The relative taxonomic composition was normalized as described above. The similarity search algorithm BLAST was performed on the three SSR viromes against the twenty viromes obtained in MetaVir. The distance matrix of taxonomic composition with relative abundance was used in a nMDS analysis to plot viromes (the metaMDS function with a Bray-Curtis dissimilarity index of the package VEGAN from R software(71)) and a PERMANOVA test (P-test) was also performed.

### Metagenomic assembly and function analysis

SSR virome assemblies were performed via a random subsampling approach as previously described(72), designed to obtain as long as possible contigs by reducing the microdiversity within the samples(73, 74). Briefly, the assembling strategy was based on random selection of a subset of the reads: 1% (75*), 5% (50*), 10% (50*), 25% (25*), 75% (25*), 100% (1*) from each sample and then assembling these subsets individually with IDBA_UD (v 1.1.2)(75) using the default parameters. Combing contigs derived from all the assemblies of the same samples and removing those < 500 bp. To this end, contigs were clustered at 90% global average nucleotide identity with cd-hit-est (v 4.7, options: −c 0.9 −n 8)(76). The relative abundance of each non-redundancy(nr) contigs was determined based on the mapping of the quality-filtered reads to the contigs, computed with bowtie2(v 2.3.3.1)(77) and SAMtools (78), using the default parameters (the total length of reads mapping to the contig divided by the contig length). Then the nr contigs with a relative abundance were uploaded to the IMG system (https://img.jgi.doe.gov/)(79), and analyzed with the standard operating procedure of the DOE-JGI Metagenome Annotation Pipeline (MAP v.4)(80). Finally, the IMG genomes are 3300028548, 3300028550 and 3300028925 respectively, were obtained. The functional content was further characterized using MG-RAST (81)(with MG-RAST accession number 4808192.3, 4808195.3 and 4808193.3 respectively), an online metagenome annotation service (http://metagenomics.anl.gov/), which compared data to the SEED Subsystems database using a maximum E-value of 1e^−5^, a minimum identity of 60%, and a minimum alignment length of 15.

### Phylogenetic trees

Two dsDNA markers: the phage terminase large-subunit domains (TerL) was present in phages of the order *Caudovirales* [Terminase_6, PF03237], and the major capsid protein (MCP) gene was present in large eukaryotic DNA virus [Capsid_NCLDV PF04451]. Both of them were used to construct the phylogenetic trees and the TerL sequences were dereplicated at the 97% nucleotide level using cd-hit(76). These markers from the SSR virome genes were screened by the DOE-JGI Metagenome Annotation Pipeline and compared to the viral RefSeq database using BLASTP (E-value < 1e^−5^) to recruit relevant reference sequences. All sequences were aligned at the amino acid level using MUSCLE(82) (using default parameters) and both maximum likelihood(ML) trees (MCP and TerL) with 1000 bootstraps were constructed with the program FastTree (v2.1.10) (83)using a JTT+CAT model and an estimation of the gamma parameter. Finally, visualized and displayed using iTOL (Interactive Tree of Life)(84).

### Genomic comparison

The Pgvv-like genomes were annotated with RAST and predicted open reading frames (ORFs) were searched against the NCBI reference viral protein(taxid:10239) with online BLASTP(85). The partial functional annotations of the Pgvv reference sequence was obtained from Yutin et al(33). Visualization of genomes map comparisons was generated with EasyFig(86).

## Conclusion

Analysis of the SSR viromes has showed that there are novel, oceanic-related viromes. A high proportion of sequence reads was classified as unknown, with only 3.31-10.87% having known virus counterparts, among these members of the order *Caudovirales* were most abundant. This pattern is consistent with previously described viromes from the Pacific Ocean as well as from a range of different biomes. The diversity of the *Caudovirales* and NCLDV in the SSR viromes is high, suggesting that in gelid environments viral diversity is high. However, the abundance and diversity of ssDNA and RNA viruses need further research. The strong signatures of Pgvv were found in the SSR, which may indicate that the virophage play an important role in regulating virus-host interaction.

## Acknowledgements

We are grateful to the funding of the National Natural Science Foundation of China (No. 41676178, 41076088, and 31500339), National Key Research and Development Program of China (2017YFA0603200), Scientific and Technological Innovation Project Financially Supported by Qingdao National Laboratory for Marine Science and Technology (No. 2016ASKJ14), and Fundamental Research Funds for the Central University of Ocean University of China (Grant Nos. 201812002, 201762017, 201562018)

